# MilkOligoCorpus: a semantically annotated resource for knowledge extraction on mammalian milk oligosaccharides

**DOI:** 10.1101/2025.02.09.637314

**Authors:** Mathilde Rumeau, Marine Courtin, Robert Bossy, Clara Sauvion, Valentin Loux, Mouhamadou Ba, Christelle Knudsen, Sylvie Combes, Claire Nédellec, Louise Deléger

## Abstract

Milk oligosaccharides are bioactive components that regulate the composition of the neonatal microbiota and exert immunomodulatory functions. Their beneficial effects depend on their structure. Numerous studies have shown intra- and inter-species variation in the structural composition and concentration of these compounds in mammalian milk, yet the biological significance of such variation remains poorly understood. Automated natural language processing methods are promising tools for extracting and gathering structured data from unstructured texts to get insight into the biological significance of milk oligosaccharide variation across mammals. These methods require training and evaluation on manually annotated text corpora. While annotated corpora exist for chemical substances, none are specifically designed for training natural language processing models to extract information on milk oligosaccharides.

To this end, we propose MilkOligoCorpus, a new gold standard for milk oligosaccharide composition in mammalian species. MilkOligoCorpus’ annotation scheme is a rich entity/relation model designed to describe the diversity pattern of milk oligosaccharides according to female factor variability and to help better understand the structure-related function of milk oligosaccharides. MilkOligoCorpus consists of 30 PubMed texts fully annotated with entities related to individuals, samples, oligosaccharides and oligosaccharide quantification linked by binary and n-ary relationships. To address data interoperability across disparate publications and databases, four terminological resources were also developed to assign unique identifiers to the entities, supported by external ontologies.

This paper presents the creation of the MilkOligoCorpus and its associated schema, along with the development of annotation guidelines and terminological resources. We also present experimental results obtained by baseline information extraction models on the corpus.

## Introduction

### Background

As the first food available to neonates in mammalian species, milk must meet their specific needs by providing ideal nutrition as well as non-nutritional bio-active components that serve physiological functions. Among the latter, milk oligosaccharides (MOs) have recently attracted increased interest regarding the wide range of potential benefits for short- and long-term mammalian health. These carbohydrates are not digested by host enzymes and reach the lower part of the gastrointestinal tract intact, where they regulate host microbiota composition and exert immunomodulatory function [1,2]. MOs are highly complex molecules, combinations of three to twenty monosaccharides and multiple linkages resulting in structurally complex matrices of linear and branched oligosaccharides. The specific functions of MOs depend on the structural organization of their molecules. For example, MOs containing α1-2-fucose motif inhibit the adhesion of the pathogen *Campylobacter jejuni* to the mouse gut [3] and enteropathogenic *Escherichia coli* to epithelial cells [4], whereas sialylated MOs have been shown to modulate the immune response [5].

Recent advancements in MO analytical technologies have allowed the investigation of oligosaccharides in the milk of numerous mammalian species [6]. These data have revealed a wide diversity in MO patterns, i.e., the abundance and richness of milk oligosaccharides in a specific species. This has led to a growing volume of scientific literature on this topic. Efforts have been undertaken to gather and consolidate literature information on the variability of inter-specific MO profiles. To date, two databases have been developed to explore MO profiles across mammals. MilkOligoDB [7] compiles MO structural diversity across mammals through the manual harvest of MOs from the literature, while the database developed by Jin et al. [8] records MO composition in 168 mammalian species. Both are exhaustive MO databases. However, they rely on manual curation of the literature, which is not suitable for processing the growing volume of biological data encompassing thousands of articles. In addition to species-specific variability [9], MO patterns also exhibit a variability related to intra-specific factors, including genotype [10] or breed [11,12], maternal physiology, i.e., duration since the end of gestation and number of gestations [13], course of lactation within a same individual [14], as well as extrinsic factors such as nutrition [15] or geography or season [16]. Understanding the biological significance of such variations in MO profiles is of importance to understand the structure-function link of MOs. That information is not collected or included in the above-mentioned databases.

This mass of data on associated or causal variability of the MOs structure patterns, spread in an increasing and disparate volume of heterogeneous scientific articles, makes it challenging to carry out manual bibliographic queries and in-depth reading across the entire study area. Automatic Natural Language Processing (NLP) methods - especially methods that extract information from textual data and that allow to store them in a database for easy querying, reuse and sharing - are particularly well-suited to handle linguistic diversity and the increasing number of publications. They also assist in studying and understanding the associated or causal variability of MOs. The first step before using NLP-based information extraction methods is to build an annotation schema that translates the biological complexity of the variability of MO profiles and conceptualize the information model. A text annotation schema is designed to identify the significant entities of interest and establish relations between them. The information schema serves both to define the NLP information extraction objective and to establish the data storage model. Regarding entities, minimal information would include the quantity of a specific milk oligosaccharide structure found in the sample of a given species. Additionally, relevant information to convey this biological complexity must include all the female intrinsic and extrinsic factors influencing MO composition, such as the number of gestations the female has already had, the duration since the mother has given birth, and the breed of the female studied. Finally, an accurate and exhaustive analysis of the diversity of milk oligosaccharides identified among mammalian species must include experimental parameters since they can influence results interpretation. They include the number of individuals studied and the methodology of analysis. Milk oligosaccharide analysis can be carried out with several methods, i.e., instruments of measurement, products used to prepare samples, and techniques to analyze results. The precision of results depends on the chosen methods, resulting in variations in their interpretation. Therefore, it is recommended to clearly specify the sample analysis methodology [17]. The recognition in the document content of the relevant entities, i.e., named entity recognition (NER), their categorization with ontology classes, i.e., entity normalization (or entity linking), and the establishment of relations between them, i.e., relation extraction, to make it available in a structured and standardized format directly integrable in a database is further complicated with this disparate and large amount of information. Usual text annotation schema relationships are binary, they consist of two entities linked by a relation. However, to clearly reflect the intricacy of the information, we also need to consider relations linking more than two entities, which adds a degree of complexity.

Another important issue lies in the intricacy of MO molecules, notably the naming variations of these chemical compounds with significant typographical variants involving hyphens, brackets, Greek letters, or word order. For example, the well-known fucosylated oligosaccharides *2’-fucosyllactose* is also referred to by its abbreviation *2’-FL*, while its systemic name, following the International Union of Pure and Applied Chemistry (IUPAC) nomenclature, is *α-L-Fucopyranosyl-(1→2)-β-D-galactopyranosyl- (1→4)-D-glucose*. Moreover, the nomenclature rules of the IUPAC are not always followed by authors, and chemical conventions used may differ between articles. For instance, Gopal et al. refer to *N-acetylneuraminic acid* as *NeuAc* when describing bovine milk composition [18], while Difilippo et al. use *Neu5Ac* in their analysis of pig milk [19]. To overcome naming variation, text annotation has to rely on semantic resources and glossaries to normalize the entities to a unified semantic representation. This normalization (named entity linking) addresses the interoperability of the data across disparate publications and databases. In fact, although our primary data source is scientific literature, we acknowledge the importance of other datasets [20] where the format of the information can vary greatly.

Beside the construction of an annotation schema fitting the complexity of MO diversity pattern, the development of NLP methods in such complex cases requires training and testing on manually annotated text corpora. Annotated corpora help evaluate the ability of the information extraction methods to identify entity types (named entity recognition) and relations, and to unify entities with respect to an external reference. To the best of our knowledge, no such corpus is available in the domain of milk oligosaccharide composition.

The present work describes the development of an annotation schema that formalizes the biological problem in an entity/relation model. This schema provides a relevant and valuable basis for modeling MO diversity pattern according to intrinsic and extrinsic variability factors in females. Additionally, it contributes to a deeper understanding of the structure-related function of MOs in health. It also describes the creation of a gold standard corpus MilkOligoCorpus for training and evaluating named entity recognition and normalization methods, and relation extraction tools meant to extract MO composition from the scientific literature. Finally, it discusses the performance of baseline methods applied to the corpus to demonstrate its relevance for evaluating and training NLP methods and to highlight information extraction challenges.

### State-of-the-art on corpora and annotation models

Before undertaking the building of the corpus, we conducted a survey of existing corpora to ascertain the potential reuse of annotations for our information extraction purposes regarding the presence of oligosaccharides in mammalian milk.

#### Chemicals

The primary entity type in our study is milk oligosaccharides. Few existing corpora include annotations for chemical compounds. Krallinger et al. provide a survey of the publicly available corpora [21]. The CRAFT (*Colorado Richly Annotated Full Text*) corpus includes annotations of compounds with ChEBI (*Chemical Entities of Biological Interest*) concepts and biological taxa with the NCBI (*National Center for Biotechnology Information*) taxonomy [22]. However, the chemical type is described with a level of precision (e.g., subatomic particles, atoms, molecules) that is not suitable for our purposes, and the CRAFT sources were selected according to the *Mouse Genome Informatics* (MGI) group, which is not relevant to our milk analysis study. Other corpora are either unavailable or out of scope (e.g., GENIA [23], ChemEVAL [24]).

The ChemDNER corpus [21] was designed to provide extensive coverage of chemical compounds and drugs in the literature. The documents were selected from PubMed using general keywords. Its annotations include over 84,000 manually identified chemical mentions, corresponding to nearly 20,000 unique chemical name strings normalized by IUPAC (*International Union of Pure and Applied Chemistry*) nomenclature and InChI (*International Chemical Identifier*) for formulas. However, despite this broad coverage, the corpus contains very few mentions of oligosaccharides, and their sample types are not documented.

#### Species

Most corpora with species annotations are related to biomedicine, focusing on human, animal models (e.g., mice, rats) and microbes. The S1000 corpus is dedicated to biodiversity surveys [25], with its protist subset containing 671 scientific articles and 1104 species annotations ranging from green plants to fungi. The low number of mammalian mentions and the absence of chemical annotations in S1000 render it irrelevant for our purposes. The *Bacteria Biotope* corpus [26] is dedicated to bacteria habitats and therefore contains many mentions of mammals related to positive or negative microbial flora. However, the mammalian mentions are not tagged with NCBI taxa, as required by our study.

#### Geographical locations

The geographical location information holds minor importance for mammalian oligosaccharide literature. The relevance of geographical location stems from its linkage to mammalian data, a feature absents in any existing corpus. We consider ChemDNER, S1000, and Bacteria Biotope relevant for evaluating the generality of the named-entity methods but inappropriate for training in our specific study domain.

### Relationships

The relationships among chemicals, media, species, their physiological stages, and locations are specific to our research topic. Most corpora in the biology domain are heavily focused on human health, where such information has not been annotated. The absence of a relevant corpus for text mining studies on milk oligosaccharides motivated our work.

## Materials and methods

This section details the process of building MilkOligoCorpus, the gold standard corpus on milk oligosaccharides. It describes the selection of literature and external semantic resources for entity normalization and the annotation process. It also explains the information extraction methods selected to assess the suitability of the corpus and serve as baselines for the development of more advanced methods.

### Document selection

We formulated a query to retrieve articles about milk oligosaccharides from the PubMed (https://pubmed.ncbi.nlm.nih.gov/) and WOS (https://www.webofscience.com/wos) bibliographic databases as they contain an extensive collection of fully open biological literature and the indexation make search efficient. The documents of MilkOligoCorpus were selected using the query in Table 1.

**Table 1:** PubMed query for the initial selection of documents (October 2022).

| # | Search Query | Results |
| --- | --- | --- |
| 1 | TI = (oligosaccharide* OR glycan*) | 28720 |
| 2 | AB = (oligosaccharide* OR glycan*) | 58887 |
| 3 | AK = (oligosaccharide* OR glycan*) | 15729 |
| 4 | #1 OR #2 OR #3 | 73135 |
| 5 | TI = (milk OR "breast feed*") | 120167 |
| 6 | AB = (milk OR "breast feed*") | 169285 |
| 7 | AK = (milk OR "breast feed*") | 58499 |
| 8 | #5 OR #6 OR #7 | 227381 |
| 9 | TI = (quantification OR composition OR content* OR concentration* OR structure*) | 2519746 |
| 10 | AB = (quantification OR composition OR content* OR concentration* OR structure*) | 10797899 |
| 11 | AK = (quantification OR composition OR content* OR concentration* OR structure*) | 969925 |
| 12 | #10 OR #11 OR #12 | 12111944 |
| 13 | #13 AND #9 AND #4 | 1602 |
| 14 | ALL = (fructose OR galacto-oligo* OR galactooligo* OR yogurt* OR cheese OR whey<br>OR FODMAPS* OR fructo-oligo* OR fructooligo* OR "enzymatic synthesis" OR<br>"enzymatic production" OR BSSL OR "nutritional yeast") | 144993 |
| 15 | TI = ("Infant formula*" OR galactosidase* OR "skim* milk" OR supplementation) | 78829 |
| 16 | ((#14) NOT #15) NOT #16 | 1161 |
| 17 | ((#14) NOT #15) NOT #16) AND Article OR Review Article OR Proceeding Paper<br>OR Early Access (Document Types) | 1134 |
| 18 | ((#14) NOT #15) NOT #16) AND Article OR Review Article OR Proceeding Paper<br>OR Early Access (Document Types) AND English (Languages) | 1108 |

The first part of the query (lines 1 to 14) selects documents about milk oligosaccharides composition. Keywords were searched within the title (TI), abstract (AB) and author keywords (AK) to maximize the likelihood of capturing a diverse range of sources, including texts where the primary focus may not be the investigation of milk oligosaccharide composition. The Glycan keyword in the query was searched as a synonym of oligosaccharide. Keywords in line 10 were used to include descriptions of all types of milk oligosaccharides, which vary from simple lists of present structures to quantification.

The queries from lines 14 to 16 helped refine our research by excluding:

● Documents studying milk oligosaccharides in non-milk samples, such as yogurt or cheese. We target analysis of raw milk as any processing or intervention can potentially alter its composition;
● Documents about milk oligosaccharides but dedicated to other purposes such as the study of methods for synthesizing milk oligosaccharides;
● Documents with a broader study of carbohydrates, including sugar structures that do not belong to milk oligosaccharides.

No restriction in the publication timeframe was made to broaden our chance to obtain diverse articles. We, however, only considered certain types of documents: article, review article, proceeding paper or early access (document types). Additionally, we only included documents in English to avoid multilingual issues.

The query (18) was carried out in October 2022 and retrieved 1108 documents. A subset of 20 texts was manually selected for manual annotation as representative of a range of mammalian species and as covering the heterogeneity of writing styles and distinctive features. For instance, Albrecht et al. list the milk oligosaccharides discovered in a table [27], while Rostami et al. detail the findings directly in the full text [28]. Salcedo et al. mention the oligosaccharides by their composition [29], Urashima et al. refer to them by their structure [30], while Difilippo et al. use their names [19].

Because manual annotation is a time-consuming task, we decided not to annotate the full articles but to select paragraphs and sentences that were informative and included the most intricate examples. The 20 texts were divided into 30 documents belonging to two types: references (titles and abstracts) and extracts from full-text articles. Extracts have variable lengths, from a single to a few paragraphs.

### Semantic resources

Our primary data source is scientific literature but multiple datasets and standards gather major information [7,8,20,31]. Entity normalization facilitates the integration of information from heterogeneous sources by mapping entity mentions to standard categories from references when relevant, and assigning these mentions to the unique identifiers of these categories. This process gives a unified format for all entity types and enables users to perform queries based on a standard form independently of their name diversity. Therefore, we selected four external semantic resources and built four new resources for normalization (the latter are described in the Results section). The existing resources are: i) the *NCBI taxonomy* for *species* [32], ii) *Geonames* for *geographical locations* [33], iii) the *Livestock Breed Ontology* for *breeds* [34], and iv) *MilkOligoThesaurus* for *milk oligosaccharides names* [35].

The *NCBI taxonomy* and *Geonames*, frequently used for data indexing, cover our needs with their extended referencing. Regarding the *Livestock breed ontology*, although it covers a limited subset of species (buffalo, cattle, chicken, goat, horse, pig and sheep breeds) and breeds within one species, it remains the most comprehensive lexical resource for mammalian breeds available to date. Finally, the main issues in exploring literature to extract relevant information on milk oligosaccharide structure relate to the heterogeneity in the way authors refer to these molecules. Thus, normalization is highly required but no existing ontology was suitable for our task. Therefore, in preliminary work, we created the *MilkOligoThesaurus* [35] that we use here for milk oligosaccharide normalization. It gathers 245 unique oligosaccharide structures and includes a standardized name, associated with several information that can be extracted from the scientific literature, such as common synonyms and abbreviations, the monosaccharide composition, its formula in condensed form, and in abbreviated condensed form, the abbreviated systematic name, the systematic name, and the isomer group. *MilkOligoThesaurus* is accessible on the AgroPortal ontology repository https://agroportal.lirmm.fr/ontologies/MILKOLIGO/ and on the *data.gouv.fr* repository https://entrepot.recherche.data.gouv.fr/dataset.xhtml?persistentId=doi:10.57745/RA5DAC in tabular, rdf and ttl format.

### Annotation process

#### Annotation guidelines

The annotation guidelines provide the detailed instructions on how to perform text annotations. The guidelines document gives a definition of each entity, entity boundary criteria, and annotation rules, i.e., what should be tagged or not, and how to assign tagged elements to semantic resources. A wide range of examples is also depicted in the document to illustrate how to handle ambiguities and inconsistencies that may lead to annotation errors and discrepancies.

To ensure accessibility for individuals with diverse backgrounds, the document includes detailed explanations of each entity type. These guidelines are intended not only for annotators of the MilkOligoCorpus but also for adaptation and use for annotating other corpora or designing instructions in prompting methods.

The annotation lead, skilled in biological science, worked alongside with text and data mining experts to develop these guidelines essential for ensuring consistent annotation. The initial draft of guidelines, authored by a single individual, was refined through an iterative annotation process after being independently reviewed by four others.

The entire guidelines, with detailed instructions and extensive list of examples are provided in the repository (https://doi.org/10.17180/94mt-v661).

#### Annotators

Two annotators with a background in biology (a PhD student expert in milk oligosaccharides, and a BS graduate with experience in manual annotation) performed the annotation. Consistency checking was performed by an NLP researcher expert in manual annotation curation.

#### Annotation process

Annotation was conducted in three stages. In the first stage, the annotator expert in milk oligosaccharides annotated all entities and binary relations from the corpus. In the second stage, the second annotator reviewed the annotations and added normalization annotations (linking entities to entries from the semantic references) as well as n-ary relations (grouping binary relations describing the same event). Finally, we applied automatic scripts to check annotation consistency and flag potential errors, which were then reviewed manually by the NLP expert.

Detected errors included missing mentions (mentions annotated in some documents but not in others), type mismatches (identical mentions tagged with different entity types), and missing or erroneous normalization annotations (with no or unknown semantic reference identifiers).

#### Annotation tools

The annotators used two different tools to perform the annotation. The first annotator used the *tagtog* tool [36] (https://docs.tagtog.com/), which is easy to set up and use, and so more suitable for first-time annotators. The second annotator and the NLP reviewer both had experience in manual annotation and used the AlvisAE tool [37], which provides advanced functionalities such as n-ary relation annotation and entity normalization.

#### Information extraction methods

We designed the MilkOligoCorpus to provide reference data to train and evaluate automatic information systems. In order to demonstrate the corpus suitability for this purpose, we evaluated baseline methods through three main information extraction steps: entity recognition, entity normalization (also called entity linking), and relation extraction. As methods, we chose common approaches based on lexicons and rules as well as recent machine-learning techniques. The purpose was to establish baseline performance, evaluate task difficulty and assess the usability of the corpus.

#### Entity recognition

Entity recognition aims to identify textual mentions of given types relevant to a given domain. Thus, here, entities are related to the topic of mammalian milk oligosaccharides. We designed two baseline models to recognize the entities defined in MilkOligoCorpus: a simple rule-based baseline and a machine-learning (ML) -based baseline.

The rule-based approach leverages semantic resources, such as lexicons and thesaurus, and a set of simple regular expressions. It is implemented using the AlvisNLP corpus processing engine (https://github.com/Bibliome/alvisnlp). A basic preprocessing pipeline based on the Stanza natural language analysis package (https://stanfordnlp.github.io/stanza/) is also used for sentence segmentation, tokenization, lemmatization and general domain entity recognition (e.g., times, cardinals…).

A common ML-based approach to recognizing entities is to fine-tune an existing transformer-based language model to classify tokens into the target entity types. Here, we use the BioBERT cased model [38], a model specific to the biomedical domain. We fine-tuned BioBERT on MilkOligoCorpus to classify all entity types at once, in a 5-fold cross-validation setting with 5 random initializations. For geographical locations, which are common in both general-domain and specialized-domain corpora, we also evaluated an existing model from the Stanza package (the English-language NERProcessor, https://stanfordnlp.github.io/stanza/ner.html) trained to recognize general-domain entities (including locations), in addition to fine-tuning our own model.

The code for our experiments is released at https://forgemia.inra.fr/holooligo/rb-pipeline and https://forgemia.inra.fr/holooligo/ner_holooligo.

#### Entity normalization

Entity normalization consists in aligning textual mentions of entities within a corpus with their corresponding entries (i.e., class, category) in a knowledge base or lexicon. Biomedical entities have the particularity that many of them can be referred to by different textual forms (e.g., both "Triose C" and "β6’-Galactosyllactose" refer to the same oligosaccharide) while similar textual forms can correspond to different entities. It can therefore be challenging to accurately map the textual mention to the correct class. We chose BioSyn as machine-learning baseline for entity normalization because it achieved good performances on similar tasks. The BioSyn framework [39] was developed to address this challenge. BioSyn relies on a pre-trained language model that is fine-tuned using synonym marginalization, a strategy that incorporates synonym sets during training to learn robust representations for entity linking. We also implemented a lexicon-based baseline that performs simple string matching between entity mentions in the text and the labels of the entries from the semantic references with which these entities have to be normalized. The code for our experiments is released in the following repositories: https://forgemia.inra.fr/holooligo/rb-pipeline, https://forgemia.inra.fr/bibliome/geonorm, and https://forgemia.inra.fr/holooligo/nen_holooligo.

#### Relation extraction

Relation extraction is the task of detecting relationships between given entities in a text and classifying them. Relation types and argument types for these relations are predefined according to the domain. Relations may involve two entities (binary relations) or more than two entities (n-ary relations). From an information extraction viewpoint, binary relation extraction is a more common and less challenging task than n-ary relation extraction. For our baseline experiments, we only tackled the task of oriented binary relation extraction.

As is typical for this task, we look at the problem of relation extraction as a sentence classification task and use a machine-learning baseline. We augment the original text with special characters that delineate the boundaries of the reference named entities and predict whether the delineated entities are in relation with each other. We use REBERT (https://forgemia.inra.fr/bibliome/re-bert), a tool that fine-tunes various language models specifically for relation extraction. We use the BioBERT model as the foundation model, as for our entity recognition baseline. The code for our experiments is released at https://forgemia.inra.fr/holooligo/brel_holooligo.

The following sentence from Rostami et al. [28] gives an example of the input given to the relation extraction method. There are two entities, the oligosaccharide name mention "3’SL" and the sample type "milk": [Example 1] “*<< 3’SL >> was found to be by far the most prominent dog @@ milk @@ oligosaccharide*.”

The relation extraction task is to predict whether there is a relation between the two entities "3’SL" and "milk". When the annotation schema only defines one possible relation type for every pair of compatible entities, then the relation type can be deduced from the argument types, and thus there is no need to specify the type of the relation. From Example 1, the only possible relation between an *oligosaccharide name* and a *sample type* is the relation *composed of* (see the Results section below for a list of all relation types). This means that if the method predicts the correct label, "yes, there is a relation", then the system will be able to deduce from the entity types that the relation type is *composed of*.

## Results

In this section, we present our annotation schema, the statistics of the annotated corpus, and the experimental results obtained with the information extraction methods.

### Annotation schema

We developed an annotation schema including entities and relations between entities. Semantic resources associated with some entities are proposed for concept normalization. The relations include binary and n-ary relations. The schema was built through an iterative process while performing the annotation tasks presented in the annotation process part of this article. The full version of the schema is the result of restructuring steps in the course of the annotation process (Fig. 1).

**Fig. 1:**
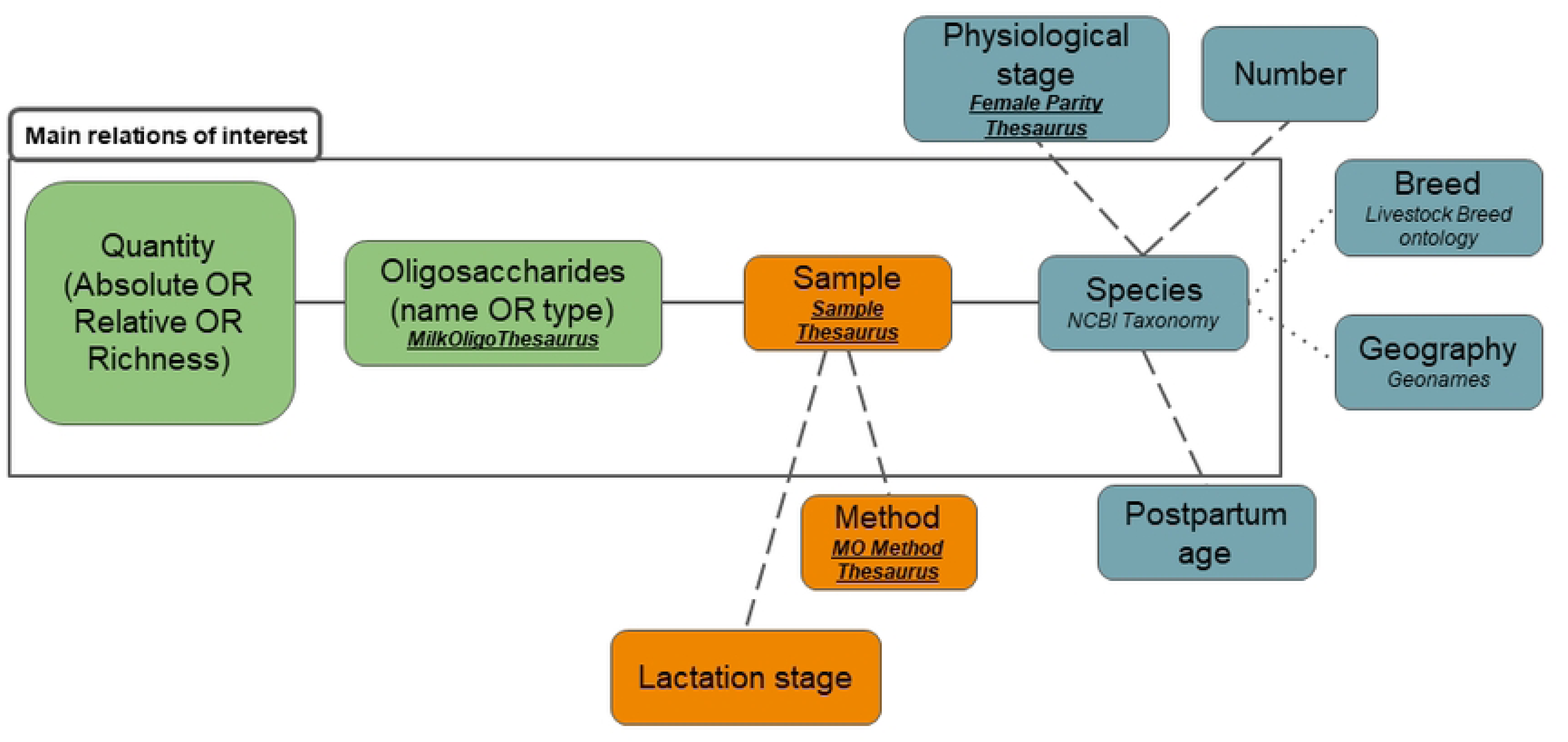
Simplified version of the annotation schema. Colors indicate the classification of entities in the three themes described above, semantic resources are in italic with proprietary glossaries in bold and underlined. Solid lines indicate priorities relations, dash lines are used for relations of minor priorities, dot lines represent the lowest-priority relations.

### Entities

Studies on mammalian milk composition include experiment records such as the number of individuals sampled, the geographical location where the sampling took place, the sampling methods, and the procedures and devices used to analyze the samples. Some studies give detailed metadata on female samples, namely breed, stage of lactation and number of gestations (primiparous, multiparous). Initially, the information included only six entity types: animals, characteristics (i.e., lactation stage, parity or age), oligosaccharides, geographical location, sample, and methodology. Several entity types have been trimmed down by removal or merging, while others have been separated into distinct entities to cover a wide range of relevant information while avoiding ambiguities with clearly distinct entities.

Originally, *species* mentions were included in a broader entity type, “animals”, including breeds when specified. We decided to introduce a separate *breed* entity type. *Breed* is mainly referred to in articles dealing with domesticated species (farm or domestic animal). The *Livestock Breed Ontology*, presented in the methodology, was the only lexica for breed terms, however, it is still limited to certain species (buffalo, cattle, chicken, goat, horse, pig and sheep breeds). We thus restricted the annotation of the breed type to domesticated species.

Similarly, female metadata were initially gathered in a large entity type including all characteristics. A first split resulted in the creation of the entity type *female physiological stage* along with an entity type for the age of the female. The physiological stage indicates how many times the female has already been in gestation, and can significantly impact the MO concentration [13]. After several annotations we decided to create two distinct entity types, *lactation stage* and *postpartum age* instead of a unique type for the age to avoid ambiguity. While *postpartum age* indicates a clear period of time, the entity *lactation stage* is a specific step during lactation. The exact duration depends on the species studied; colostrum last only 48h for rabbits while it extends up to 30 days in giant pandas [40].

Finally, milk oligosaccharides are complex structures of core molecules composed of three monosaccharide building blocks that can be repeated up to 25 times but also decorated with two other monosaccharides. They are typically classified in three categories according to this structure, *neutral* if the structure is only composed of the core molecules and *sialylated* or *fucosylated* depending on the monosaccharide decorative. Two other categories are also used, based on the chemical linkages between the monosaccharides, *type I* or *type II*. The precision of milk oligosaccharide composition varies depending on the methodology used. Some analyses are used to identify as many structures as possible, e.g. shotgun-MS analysis, while others cannot distinguish between isomers - molecules with the same chemical composition and mass but different arrangements of atoms - such as MALDI-TOF-MS [17]. As a result, several studies have been unable to clearly identify specific molecules but have instead detected characteristics elements of these molecules, classifying them into broader categories. Given this, we have decided to add a distinct entity type for oligosaccharide types.

The methodology also influences quantitative characterization. Initially gathered in one entity, we decided to create three milk oligosaccharide quantification entities. The *absolute quantification* includes indication regarding the total concentration of oligosaccharide, while the *relative quantification* is the proportion of an oligosaccharide compared to the total milk oligosaccharide. Finally, the *richness* gives the number of an oligosaccharide or a group of milk oligosaccharides.

In the end, the annotation schema consists of fourteen entity types that we classified into three groups based on the aspect they refer to: (i) Individual-related; (ii) Sample-related; (iii) Oligosaccharide-related.

The complete list of entities is available in Table 2, along with the semantic resources described in the following paragraph.

**Table 2:** Description of the 14 entities of the annotation schema. Semantic normalization refers to semantic resources used to normalize the entities. “Indiv.-related” stands for “individual-related” and “Oligo-related” stands for “Oligosaccharide-related”.

| Entity group | Entity type | Definition | Semantic normalization | Text examples |
| --- | --- | --- | --- | --- |
| Indiv.-related | Species | Name of mammalian species (scientific name, current name, abbreviations ...) | NCBI | Porcine, dogs, mongoose lemur, bovine ... |
| Indiv.-related | Breed | Subclass of species | Livestock breed ontology | Dutch Landrace, Labrador Retriever ... |
| Indiv.-related | Geography | Indicates where the samples are from | Geonames | Cork, County Tipperary, Washington DC ... |
| Indiv.-related | Female physiological stage | Lactation parity that indicates the number of gestation mothers have already had | Female parity thesaurus | Gilt, bred at least once ... |
| Indiv.-related | Individual number | Number of individuals sampled in letter or numeric forms | No | 335, Three ... |
| Sample-related | Sample type | Refers to the samples used during the experimentation | Sample thesaurus | Milk, pooled, mixed ... |
| Sample-related | Lactation stage | Type of milk based on the period of lactation | No | Mature, early lactation, colostrum ... |
| Indiv.-related | Postpartum age | Duration in number since the mother has given birth | Number to convert in days | Day 10, 96 days postpartum ... |
| Sample-related | Methodology of analysis | Procedures and devices used to analyze the samples | MO methods thesaurus | HPAEC-PAD, NMR, Q-TOF-MS ... |
| Oligo-related | Oligosaccharide type | Neutral, fucosylated, sialylated, acidic, type I, type II | Oligo type thesaurus | Fucosylated, type II, Sia-MOS ... |
| Oligo-related | Oligosaccharide name | One of the Oligosaccharide's thesaurus mentions | MilkOligoThesaurus | 2'-Fucosyllactose, LNFP V, Gal(b1 – 3)Gal(b1 – 4)Glc ... |
| Oligo-related | Absolute quantification | Concentration of an MO (Number in g.L) | No | 30 mg/L, 19.3 +/- 2.9 g/L ... |
| Oligo-related | Relative quantification | Proportion of an MO (Number in %) | No | 2.5 %, 20-52 % ... |
| Oligo-related | Oligosaccharide richness | Number of all the MO, a single MO or a group of MOs | No | Eight, 29 ... |

### Semantic resources

We used semantic resources to normalize the relevant entities. As described in the Material and Method section, we identified existing semantic resources to normalize *species*, *geographical locations*, *breeds*, and *milk oligosaccharide names*. Additionally, we also developed our own resources to normalize the entities for which no suitable existing resource was found, i.e., *female physiological stage*, *sample type*, *methodology of analysis* and *oligosaccharide type* (Table 2). We created four glossaries, the *Female physiological stage thesaurus*, the *Sample thesaurus*, the *MO methods thesaurus* and the *Oligo type concepts*, to overcome the lack of reference sources for the *female physiological stage*, the *type of sample* studied, the *methodology* used for the analysis, and the *oligosaccharide type* respectively.

The *Female physiological stage thesaurus* gathers terms to designate the physiological stage of the female i.e., the number of gestation of females. Primiparous and multiparous are common names for parity, but we also found phrasal forms (“Bred for the first time”) or the number of parity (“x”-parity). Besides, porcine species have specific words, synonyms for pig, that indicate the parity. Sows are multiparous pigs, whereas gilts are primiparous pigs. These two physiological stages, primiparous and multiparous, are compiled in a 6-column tabular file including an identifier for each stage, synonyms, and specific characteristics relevant to the pig species. The *Female physiological stage thesaurus* is accessible in the repository https://doi.org/10.57745/LFXGFO.

This study focuses on the analysis of milk in its raw form as any intervention can potentially alter its composition. However, in some experiments, it is necessary to combine samples from multiple individuals. This is usually specified in the article and must be considered when comparing results across different experimental designs. Given the variability in formulation (adverbs, verbs, nouns) we decided to standardize it in the *Sample thesaurus*. The two types of samples, milk or pooled milk, are compiled in a 7-column tabular file that includes the identifier of the type of sample, and synonyms. The *Sample thesaurus* is accessible in the repository https://doi.org/10.57745/LFXGFO.

A third thesaurus is used to normalize the textual mentions of analysis methodology. The *Chemical Method Ontology* (CHMO) lists instruments and methods to collect data and prepare material in chemical experiments. It is an accurate ontology but it is too broad for our field of interest. Milk oligosaccharides analysis relies on a few specific methodologies which we have compiled into a concise thesaurus called the *MO methods thesaurus*. The forty-two methodology types are compiled in a 6- column tabular file that associates to each methodology an identifier, abbreviation, synonyms, the doi from the article where the methodology was mentioned, and the identifier from the CHMO database when available to enable data linking. The *MO methods thesaurus* is accessible in the repository https://doi.org/10.57745/LFXGFO.

The oligosaccharides’ types refer to the usual classification used to categorize milk oligosaccharides. This classification is based on the structure of the molecules. The *Oligosaccharide type concepts* lists the terms used and *sialylated* is the only type that can be referred to with a different name, *acidic.* The *Oligosaccharide type concepts* is accessible in the repository https://doi.org/10.57745/LFXGFO.

Finally, entities referring to quantification, i.e., *individuals’ number*, *absolute quantification*, *relative quantification*, and *oligosaccharide richness*, do not require semantic resources for normalization. Those numerical entities must only have homogenized units of measure.

### Relations

Information extraction aims to extract relevant knowledge from unstructured textual data to make them available in a structured format. To associate the MO structure pattern, i.e., the abundance and richness of milk oligosaccharides in a specific species, isolated entities alone are insufficient. It is necessary to define the relationships between them to accurately describe their association. The following sentence from Gopal et al. [18] gives an example [Example 2] of entities in relations, where the *relative quantification* entity, 50 %, is meaningful only when linked to the oligosaccharides it described, which in turn are relevant only when linked to the *species* in which they are found.

[Example 2]: *“Together, 3’- and 6’-sialyllactose account for more than 50 % of the total oligosaccharides present in bovine”*

Relations can involve two entities (binary relations) or multiple ones (n-ary relations). In our annotation schema, we defined eleven types of binary relationships, that are listed in Table 3 with the types of the arguments involved. Examples for each relation type are available in the guidelines document.

**Table 3:** Characteristics of the binary relations with the definition and argument entity types.

| Relation | Argument 1 | Argument 2 | Definition |
| --- | --- | --- | --- |
| Is a | Species | Breed | Relates the species to its breed |
| From | Species | Geography | Relates the species to its location |
|  | Sample Type | Geography | Relates the sample to its location |
|  | Lactation Stage | Geography | Relates the lactation stage to its location |
| Number of | Breed | Individual Number | Relates the breed to the number of individuals |
|  | Species | Individual Number | Relates the species to the number of individuals |
| Has a physiological stage | Species | Female<br>Physiological Stage | Relates the species to its physiological stage |
| Has given birth since | Species | Postpartum Age | Relates the species to the period since when she has given birth |
| Has produced | Species | Sample Type | Relates the species with the milk |
|  | Species | Lactation Stage | Relates the species with the milk |
| Collected at the stage | Sample Type | Lactation Stage | Indicates the stage of lactation at the time of sampling |
| Collected x days after parturition | Lactation Stage | Postpartum Age | Indicates the duration between the parturition and the collection of the samples |
|  | Sample Type | Postpartum Age | Indicates the duration between the parturition and the collection of the samples |
| Analyzed by | Lactation Stage | Methodology of analysis | Indicates the methodology used to analyze the samples |
|  | Sample Type | Methodology of analysis | Indicates the methodology used to analyze the samples |
| Composed of | Sample type | Oligosaccharide name | Indicates the type of oligosaccharide found in a sample |
|  | Lactation stage | Oligosaccharide name | Indicates the type of oligosaccharide found during a specific lactation stage |
|  | Sample type | Oligosaccharide name | Indicates the oligosaccharide found in a sample |
|  | Lactation stage | Oligosaccharide name | Indicates the oligosaccharide found during a specific lactation stage |
| Found in quantity | Oligosaccharide type | Absolute quantification | Indicates the absolute quantity of a type of oligosaccharide |
|  | Oligosaccharide type | Relative quantification | Indicates the relative quantity of a type of oligosaccharide |
|  | Oligosaccharide type | Oligosaccharide richness | Indicates the number of a type of oligosaccharide |
|  | Oligosaccharide name | Absolute quantification | Indicates the absolute quantity of an oligosaccharide |
|  | Oligosaccharide name | Relative quantification | Indicates the relative quantity of an oligosaccharide |
|  | Oligosaccharide name | Oligosaccharide richness | Indicates the number of an oligosaccharide |

Given the definitions of the relationships *from*, *composed of* and *found in quantity*, the argument types are interchangeable. The *from* relation can connect a *geography* entity either to *species*, *sample type* or *lactation stage* entities, depending on the context. For relations involving milk oligosaccharide entities (*composed of* and *found in quantity*), the variation arises from differences in the precision of the molecule details i.e., whether the study specifies the exact molecule names or only the type, and in quantification, distinguishing between the exact number of molecules or a relative quantity.

Regarding the relations *number of*, *has produced*, *collected x days after parturition* and *analyzed by* we decided to select interchangeable argument types based on the distance between arguments. The arguments to include in the relations were chosen after consideration of the location of the entity mentioned in the text. Relation extraction (RE) methods typically give better performance when entities are close by in the text. However, critical relations may occur between entities mentioned in separate and sometimes very distant sentences. In fact, to avoid redundancy authors do not always repeat the information already stated. For instance, in Salcedo et al. [29] the relation *composed of* links “milk” and “fucosylated” located in two distinct sentences [Example 3]: [Example 3] “*<u>Porcine</u> <u>milk</u> had lower OS diversity […] In agreement with previous studies, only <u>3 fucosylated</u> OS were identified*.”

Similarly, in Urashima et al. [41] the relation *analyzed by* includes the *methodology of analysis* (“gel filtration and preparative thin layer chromatography”) and the *sample studied* (“milk”) which is mentioned in the previous sentence [Example 4]:

[Example 4] “*Carbohydrates were extracted from high <u>Arctic harbour seal milk</u>, Phoca vitulina vitulina (family Phocidae). Free <u>neutral</u> oligosaccharides were separated by <u>gel filtration and preparative thin layer chromatography</u> […].*”

We chose to annotate cross-sentence relations between entities occurring in the same paragraph and not further away. However, to simplify the task for very distant entities, we decided to include flexible relations where one argument type may be replaced by another one close in meaning, such as in the *has produced* relation where the entity type *lactation stage* can be used instead of the entity type s*ample type*. This flexibility does not provoke disparities.

Furthermore, considering the number of entities to be extracted, binary relations may not be sufficient to capture the information unlike n-ary relations that link more than two entities. The sentence in example 5 gives an example of MO (referred to as OS in example 5) quantities measured in the milk of different species where four arguments must be linked to fully capture the meaning [42].

[Example 5]: “*Approximately 29 different OS were identified in porcine milk, compared to 40 in bovine milk and >130 in human milk.”*

In this sentence, we need to establish three 4-ary relations indicating the quantity of milk oligosaccharides in the species: (i) 29 MOs in porcine milk; (ii) 40 MOs in bovine milk; (iii) 130 MOs in human milk.

Since, “OS” is stated only once, a relation extraction method relying solely on binary relations cannot accurately determine the number of MOs present in the different species. It would link all numbers, 29, 40 and >130 to OS, the latter being related to milk, itself linked to porcine, bovine or human. We cannot simply merge the binary relations because it would not indicate the number of MOs associated with the respective species.

N-ary relations are thus an essential part of our annotation schema. Fig. 1 is a simplification of our annotation schema with entities and relations organized in hierarchies based on the types of lines representing the relations. We have grouped entities by theme: (i) Individual-related entities are in blue; (ii) Sample-related entities are in orange; (iii) Oligosaccharide-related entities are in green.

As the number of arguments increases, automatically extracting relations becomes even more challenging. Therefore, we have prioritized the relations to be extracted, ensuring that efforts are primarily focused on the key relations of interest. These relations are framed in **Error! Reference source not found.** by “Main relations of interest”.

The entire schema represents the maximal n-ary relation and each subgraph is a potential relation. The main relations of interest contain four entities: s*pecies*; *sample*; o*ligosaccharides* and *oligosaccharide quantities*. This 4-ary relation contains the central biological information to describe MO patterns in mammalian species. Entities linked by dash lines bring a higher level of detail.

The two 5-ary relations in the sentence below [Example 6] from Rostami et al. [28] include detailed information on the *species*, *postpartum age* or *lactation stage* which indicate relevant biological information regarding the evolution of MOs during the course of lactation within a same individual. The extraction of this information is a key to understanding the biological significance of such variations in MO profiles.

[Example 6]: *<u>3’SL</u> was found to be by far the most prominent <u>dog milk</u> oligosaccharide. In the <u>first days of lactation</u> it was present at levels around <u>7.5 g/L</u> (12 mM), which dropped rapidly to approximately <u>1.5 g/L</u> at <u>10 days of lactation</u>*.

Finally, we estimated that *geography* and *breed* represent the least critical information to extract. While they provide relevant information, these entities appear less frequently in texts and add complexity to the n-ary relation extraction task. As shown in Example 7 below from [43], a 6-ary relation, including *species*, *oligosaccharide name*, *geography*, *sample*, *postpartum age*, and *oligosaccharide absolute quantification*, is necessary to address the differences in human MO (HMO in Example 7) concentration according to the *postpartum age*.

[Example 7]: “*A decrease (P < 0.05) in HMO concentration was observed during the course of lactation for the US mothers, corresponding to 19.3 +/- 2.9 g/L for milk collected on postnatal day 10, decreasing to 8.53 +/- 1.18 g/L on day 120 (repeated measures; n = 14).”*

In many cases, n-ary relations are necessary to resolve ambiguities and extract the most detailed information possible. The annotation schema takes into account the challenge of n-ary relation extraction by prioritizing information. This prioritization does not exclude entities but instead establishes a hierarchy in cases of complex extraction aiding decision-making when necessary. We identified the key milk oligosaccharides information of scientific relevance to support the development of automatic methods which will be the focus of development and training efforts.

### Corpus

The resulting corpus is a collection of 30 documents annotated with 3626 entities, 2663 binary relations and 1927 n-ary relations. The corpus is available in the following repository: https://doi.org/10.57745/LFXGFO.

Table 4 shows the number of occurrences of entities of each type, the number of distinct surface forms, and the ratio between the number of distinct forms and the number of occurrences. This ratio is an indicator of the variability of each entity type and thus the expected difficulty to recognize them automatically. The most common entity type is *oligosaccharide name*, which is expected because this entity type is central to the MilkOligoCorpus. Nevertheless, the variability of *oligosaccharide name* entities is relatively low. This may seem surprising since molecule names tend to show a lot of variability, however, at least 254 occurrences among 913 are very generic terms, like “HMOs”, “PMOs”, or “oligosaccharides”. *Sample type* entities show an extremely low variability, indeed, 98% of occurrences have the form “milk” or a trivial variation, like “Milk”, or “milks”. The same phenomenon can be observed for *lactation stage* for which 45% of occurrences are “colostrum”, and for *female physiological stage* where 83% of occurrences are “gilt”. The most variable entities are those of types that represent numeric values: *postpartum age*, *oligosaccharide richness*, *relative quantification*, *absolute quantification*. Indeed, each distinct number is counted as a different form. On the other hand, the high variability of *geography* is due to the high number of different places mentioned in the corpus.

**Table 4:** Statistics of each entity type in the MilkOligoCorpus.

| Entity Type | Number of occurrences | Number of distinct forms | Ratio |
| --- | --- | --- | --- |
| Oligosaccharide name | 913 | 277 | 0.30 |
| Species | 754 | 100 | 0.13 |
| Sample Type | 635 | 8 | 0.01 |
| Oligosaccharide type | 293 | 68 | 0.23 |
| Lactation stage | 289 | 35 | 0.12 |
| Postpartum Age | 196 | 132 | 0.67 |
| Oligosaccharide richness | 134 | 59 | 0.44 |
| Methodology of analysis | 84 | 61 | 0.73 |
| Relative quantification | 79 | 61 | 0.77 |
| Geography | 64 | 46 | 0.72 |
| Female Physiological Stage | 59 | 8 | 0.14 |
| Individual Number | 47 | 20 | 0.43 |
| Breed | 45 | 25 | 0.56 |
| Absolute quantification | 34 | 30 | 0.88 |
|  | Total: 3626 | Average: 66.43 | Average: 0.44 |

Table 5 presents the breakdown of binary relations by type. *Composed of* and *has produced* are by far the most frequently occurring binary relations. The prevalence of these two binary relation types reflects the fact that, in the annotation schema, their arguments belong to the most common entity types: *oligosaccharide name*, *oligosaccharide type*, *species*, *sample type*.

**Table 5:**
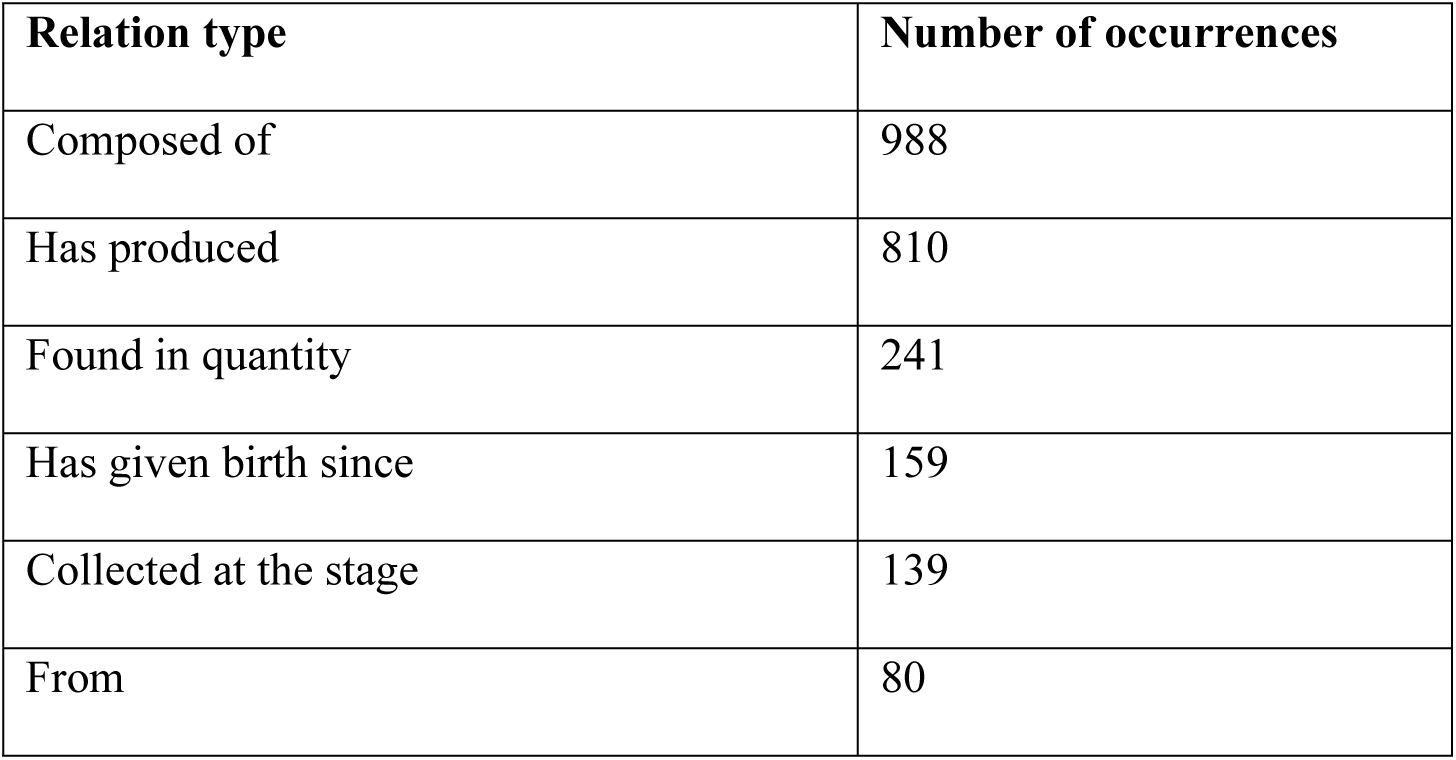
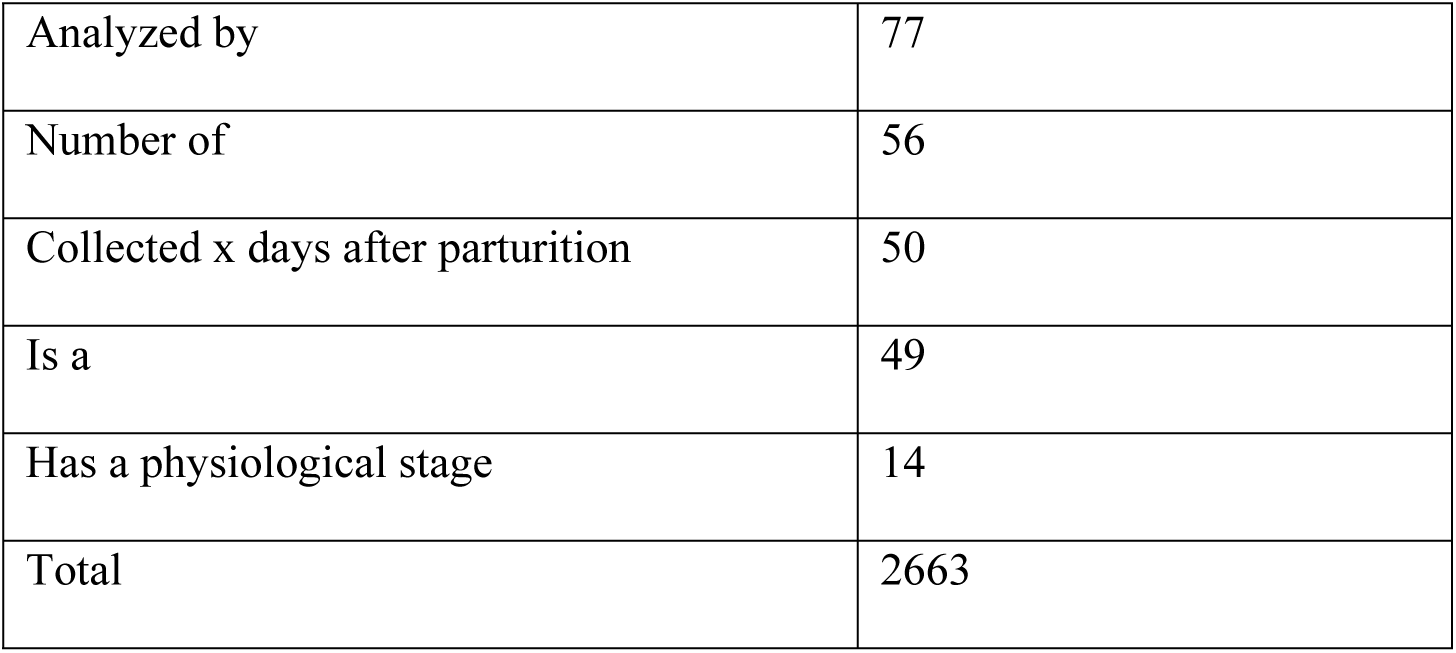
Statistics of each binary relation type in the MilkOligoCorpus.

N-ary relations are built from the aggregation of binary relations pertaining to the same knowledge or event. Table 6 shows the distribution of n-ary relations by arity. Arity is the number of arguments in the relation. An arity of 2 indicates a binary relation that has not been aggregated. Out of 2663 binary relations, 18% are not aggregated with other relations. Among the n-ary relations of arity above 2, 87% are ternary or quaternary relations. Higher arity relations are rare.

**Table 6:** Arity statistics for n-ary relations.

| Arity | Number of relations |
| --- | --- |
| 2 | 470 |
| 3 | 796 |
| 4 | 475 |
| 5 | 141 |
| 6 | 42 |
| 7 | 1 |
| Total | 1925 |

### Experimental results

We report the performance of the baseline methods on MilkOligoCorpus for the three information extraction tasks, entity recognition and normalization, and relation extraction. We use standard evaluation metrics: precision, recall and F-score for entity recognition and relation extraction, and accuracy for entity normalization.

### Entity Recognition: Rule-based methods

Table 7 shows the performance of the rule-based approach. Overall, the baseline achieves satisfactory performance, particularly for entity types such as *lactation stage* and *female physiological stage*, which are well-covered by the lexicons (Table 2). Similarly, *oligosaccharide names* and *types*, as well as mammal *species* are identified effectively. For *species*, a significant disparity is observed between precision and recall. This discrepancy stems from the absence of certain vernacular names (e.g., "sow" or "porcine") or common terms (e.g., "mothers") from the lexicon. These frequent terms account for the relatively low recall for this entity type.

**Table 7:** Comparison of F-score, precision and recall measures across entity types for rule-based named entity recognition (NER). The evaluation mode is strict, requiring exact entity boundary matching with the reference.

| Entity Type | F-score | Precision | Recall |
| --- | --- | --- | --- |
| Overall (Micro average) | 0.74 | 0.82 | 0.67 |
| Sample Type | 0.98 | 0.98 | 0.99 |
| Species | 0.73 | 0.95 | 0.59 |
| Lactation Stage | 0.82 | 0.89 | 0.75 |
| Oligosaccharide Type | 0.62 | 0.65 | 0.60 |
| Oligosaccharide Name | 0.79 | 0.85 | 0.73 |
| Postpartum Age | 0.29 | 0.53 | 0.20 |
| Oligosaccharide Richness | 0.25 | 0.68 | 0.16 |
| Individual Number | 0.38 | 0.75 | 0.26 |
| Relative Quantification | 0.45 | 0.41 | 0.52 |
| Methodology of Analysis | 0.34 | 0.31 | 0.38 |
| Breed | 0.29 | 0.31 | 0.27 |
| Absolute Quantification | 0.58 | 0.64 | 0.53 |
| Female Physiological Stage | 0.98 | 1 | 0.97 |

For *breeds*, the baseline relies on the *Livestock Breed Ontology*, which focuses exclusively on livestock species (e.g., cattle, chicken) and does not cover other mammalian species found in the corpus, which explains the low performance.

Unsurprisingly, entities with a high distinct-form-to-occurrence ratio, as shown in Table 4, tend to exhibit lower performance with this simple lexicon-based baseline. Entities involving numerical values, such as *oligosaccharide richness* or *number of individuals*, are particularly challenging to detect accurately without introducing excessive noise. This is true as well for methodologies of analysis, where the variation in terminology is not comprehensively captured by the thesaurus that we used.

It should be noted that we did not use a lexicon-based approach for *geographical* entities, since the *GeoNames* lexicon contains too many ambiguous names that may be confused with other words. For instance, *GeoNames* includes location names like “Milk” which in some rare contexts may refer to the name of a geographical place, but most likely never in our corpus. An approach relying on lexicon matching would certainly give extremely poor results, so we did not attempt it.

### Entity Recognition: Machine learning methods

The results of our fine-tuned BioBERT model are shown in Table 8. They are computed as the averages over the 25 runs (5-fold cross-validation with 5 random initializations), with the standard deviation reported as well. The main entities (*species*, *sample type*, *oligosaccharide type* and *oligosaccharide name*) are well-identified, with the exception of quantities (*relative quantification* and *absolute quantification*). *Postpartum age*, *breed* and *geographical location* types also yielded low scores. There are a number of possible explanations. Quantities are dependent on the presence of *oligosaccharide* entities, which makes it harder for the model to learn, especially when the number of examples is relatively small. *Postpartum ages* are often discontinuous entities, which are not fully handled by the model. *Geographical locations* and *breeds* are rare in the corpus and have variable forms (Table 4). For *methodology of analysis* entities, which are also infrequent, the score is average (0.57 F-score). Only the score for *physiological stage* type is extremely low. The model is unable to learn this entity type. This phenomenon can be attributed to two main reasons: the low occurrence frequency of this entity in the corpus and, most importantly, the fact that there are many instances where the same span is both of a *female physiological stage* type and a *species* type for example, "gilt" refers to a female pig (*species*) that has not yet given birth to piglets (*female physiological stage*). This case is not handled by our baseline method that tags mentions by a single type.

**Table 8:**
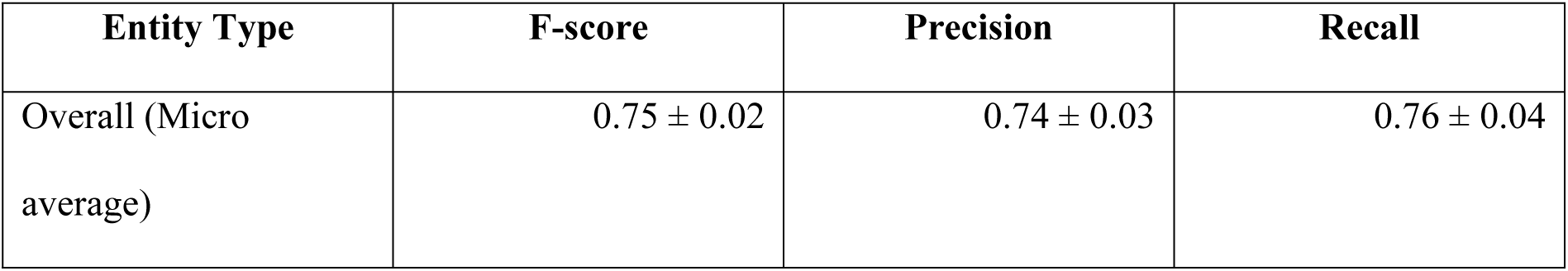
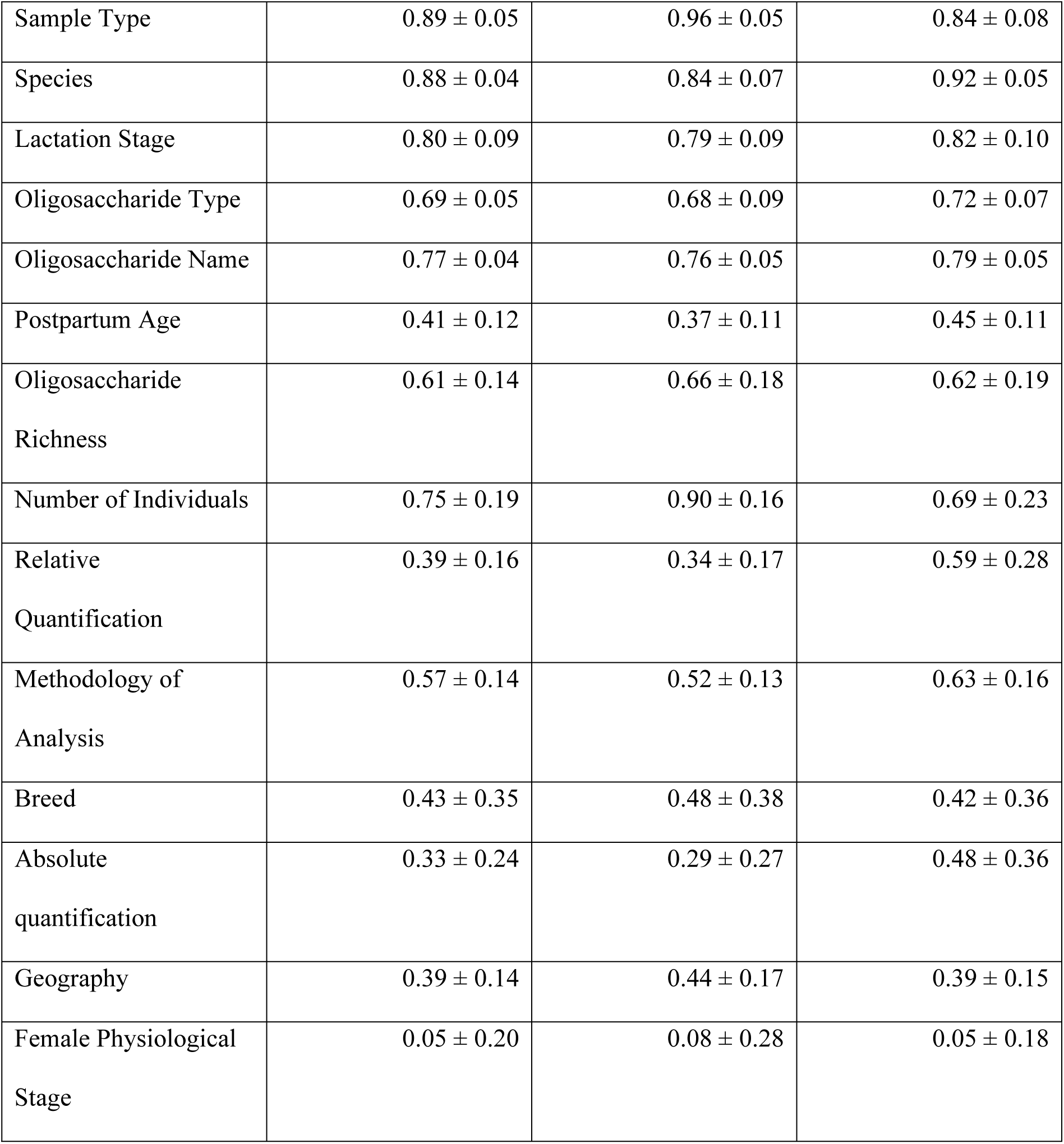
Comparison of f-score, precision and recall performance across entity types obtained with the machine-learning method. The evaluation mode is strict, requiring exact boundary matching with the reference.

As mentioned in the Methods section, we also evaluated Stanza’s model for geographical location recognition. It achieved a precision of 0.58, a recall of 0.72 and an F-score of 0.64. These scores are higher than those of our own model (F-score of 0.39, precision of 0.44 and recall of 0.39), likely due to Stanza’s model being trained on a significantly larger corpus. But they are only moderately good, largely due to differences in the training data. Stanza’s model was trained on an annotated corpus that defines *geographical* entities slightly differently and includes document types that significantly differ from those in our corpus. As a result, some corpus-specific abbreviations, such as "MO" for *milk oligosaccharide*, were misclassified as false positives by Stanza, along with proper names like the names of the publication’s authors. Additionally, Stanza’s model frequently confused *geographical* entities with *breeds*, as certain breed names incorporate geographical locations (e.g., "Alaskan husky").

Compared to the rule-based approach, the machine-learning (ML) approach generally achieved higher recall and a better precision-recall balance, emphasizing its ability to generalize and make predictions for unseen examples. The overall F-score for ML was also slightly higher (0.75 vs. 0.74). Performance varied across entity types, with the ML approach outperforming the rule-based method for most entities. However, the rule-based approach performed better for entities like *sample types* and *female physiological stages* that showed little variation and ambiguity. It also achieved higher performance for *lactation stages* and *oligosaccharide names*, although the F-scores of both methods were similar.

### Entity Normalization: Lexicon-based approach

Table 9 shows the performance of the lexicon-based approach for entity normalization. Performance is good for most entities, except for *geographical* entities. This is explained by the high ambiguity of *geographical* names (the same name may refer to different locations depending on the context) as already stated. *Oligosaccharide types*, *sample types* and *female physiological stages* obtained perfect scores due to the very small number of classes in the semantic references and the low variability of the entity forms.

**Table 9:** Entity normalization scores obtained by the lexicon-based approach.

| Entity type / Semantic reference | Match accuracy |
| --- | --- |
| Species / NCBI | 0.72 |
| Methodology of analysis / MO methods thesaurus | 0.96 |
| Oligosaccharide name / MilkOligoThesaurus | 0.62 |
| Oligosaccharide type / Oligo type concepts | 1.00 |
| Geography / GeoNames | 0.56 |
| Sample type / Sample thesaurus | 1.00 |
| Female physiological stage / Female physiological stage thesaurus | 1.00 |

### Entity Normalization: Machine-learning approach

We used the BioSyn ML approach to normalize entities related to *methodology of analysis*, *oligosaccharide names*, and *species*. In contrast, *oligosaccharide types*, *sample types* and *female physiological stages* are normalized with a very small number of concepts and show little ambiguity, so a simple rule-based approach is sufficient to obtain high performance, making more complex ML not necessary. For *geographical* locations, the semantic reference used for normalization (*GeoNames*) is so large that it becomes computationally prohibitive for BioSyn to handle, which is why we opted not to use it. Further investigation is needed to design a computationally efficient ML approach for this entity type, which we leave for future work.

The BioSyn ML results are shown in Table 10 as averages over 5 random initializations. Performance is rather good overall, and very consistent. For *species* entities, it is much higher than with the lexicon-based approach (0.85 vs. 0.72, respectively for BioSyn vs. the lexicon-based approach). It is slightly lower but still quite high (0.92 vs. 0.96) for *methodology of analysis* entities. The performance of both baselines is similar for *oligosaccharide names*.

**Table 10:** Entity normalization scores obtained by BioSyn (mean ± standard deviation)

| Entity type / Semantic reference | Match accuracy |
| --- | --- |
| Methodology of analysis / MO methods thesaurus | 0.92 ± 0.0 |
| Oligosaccharide name / MilkOligoThesaurus | $0.62 \pm 0.02$ |
| Species / NCBI | $0.85 \pm 0.0$ |

### Binary relation extraction

Table 11 shows the scores for the relation extraction baseline in a 5-fold cross-validation setting. Performance is strong, near 70 F1-score and over, for most relation types given the complexity of the task. The relation *from* obtains slightly lower performance, which is probably due to the small number of its occurrences in the corpus. Only the *collected x days after parturition* relation gives very poor results. The model seems unable to learn this relation type. This may be due to its relatively low frequency of occurrences, as well as its strong dependence on context. This relation typically occurs only when the *species* entity is not mentioned nearby in the text and when the *has given birth since* relation between the *species* and the *postpartum age* cannot be annotated. Actually, the baseline method considers only entity pairs that appear together within a maximum of two sentences as candidates for a relation. As a result, entities that are farther apart are never predicted to be in relation with each other, leading to lower scores for most relation types. This limitation particularly affects the *composed of* and *analyzed by* relations, which have a higher proportion of long-distance relations (respectively 0.26 and 0.19).

**Table 11:** Comparison of F-score, precision and recall across relation types.

| Relation Type | F1-score | Precision | Recall |
| --- | --- | --- | --- |
| Analyzed by | $0.63 \pm 0.20$ | $0.66 \pm 0.21$ | $0.64 \pm 0.22$ |
| Collected at the stage | $0.81 \pm 0.12$ | $0.89 \pm 0.08$ | $0.76 \pm 0.19$ |
| Collected x days after parturition | $0.19 \pm 0.19$ | $0.26 \pm 0.35$ | $0.35 \pm 0.43$ |
| Composed of | $0.58 \pm 0.02$ | $0.68 \pm 0.08$ | $0.51 \pm 0.08$ |
| Found in quantity | $0.73 \pm 0.05$ | $0.75 \pm 0.09$ | $0.73 \pm 0.08$ |
| From | $0.41 \pm 0.40$ | $0.42 \pm 0.40$ | $0.39 \pm 0.40$ |
| Has a physiological stage | $0.75 \pm 0.42$ | $0.71 \pm 0.41$ | $0.80 \pm 0.45$ |
| Has given birth since | $0.61 \pm 0.20$ | $0.67 \pm 0.26$ | $0.56 \pm 0.16$ |
| Has produced | $0.80 \pm 0.03$ | $0.84 \pm 0.08$ | $0.78 \pm 0.07$ |
| Is a | $0.78 \pm 0.16$ | $0.86 \pm 0.19$ | $0.76 \pm 0.24$ |
| Number of | $0.80 \pm 0.29$ | $0.83 \pm 0.16$ | $0.84 \pm 0.36$ |
| Overall (Micro average) | $0.68 \pm 0.05$ | $0.74 \pm 0.05$ | $0.64 \pm 0.06$ |

It should also be noted that for some relations, especially the less frequent ones, the score is highly dependent on the split of the corpus, as indicated by higher standard deviations.

## Discussion

This paper presents the development of an annotation schema and the associated MilkOligoCorpus, a gold standard corpus, composed of 30 documents relevant to milk oligosaccharides composition in mammalian species. The annotation schema includes fourteen entity types, each of which is normalized, with a semantic resource, to standardize the information. Three databases, database from Remoroza et al. [20], MilkOligoDB [7], and Jin et al.’s database [8], dedicated to milk oligosaccharides are already available, offering an exhaustive overview of milk oligosaccharide composition in mammalian species. However, Remoroza et al.’s database [20] does not relate milk oligosaccharides to species while MilkOligoDB [7], and Jin et al.’s database [8] do not include metadata that are crucial to elucidate further hypotheses regarding parameters influencing milk oligosaccharides composition.

The originality of the MilkOligoCorpus relies on the richness of the annotated information. Notably, the characteristics of the experimentation (number of individuals, methodology of study, pooled milk or not), are crucial for further interpretation of the data, and detailed characteristics of females allow the development of further hypotheses.

The baseline methods we implemented and evaluated on our corpus yielded satisfactory performance overall for the three information extraction tasks despite their complexity. This demonstrates the usability of our corpus for the development of automatic methods to extract information about mammalian milk oligosaccharides. There is room for improvement in the performance, since we only tested common baselines and certain entity and relation types were more challenging. We plan to investigate these issues in future work with state-of-the-art methods, as well as the more complex task of n-ary relation extraction. We also hope that the release of our corpus and baselines will foster information extraction research in this domain.

For that purpose, the number of annotated articles may be a limitation of the MilkOligoCorpus. We have decided to put our efforts into the precision and richness of the annotation rather than into the quantity, specifically regarding the complexity of chemical structure. We intend to extend this corpus with literature used to build existing databases [7] and [8].

Our final goal is to develop methods to retrieve information from the literature that we will make publicly available in a database including oligosaccharide molecules (different nomenclatures or ways of describing molecules co-exists in the scientific community, information with different levels of availability) and the relation between these molecules and phenotypic data, species mentions, stage of lactation, milk quality and maternal parameters from unstructured data. Automatic methods appear as a very promising way to alleviate the burden of the manual analysis of papers. The baseline methods we implemented give encouraging results, but need to be further improved in future work in order to achieve a good balance between automatic extraction and manual curation.

## Conclusion and future directions

Research in milk oligosaccharides has seen accelerated growth, making it challenging to maintain a detailed overview of the published literature. This work proposed an annotation schema designed to represent entities and relations modeling mammalian milk oligosaccharides and related information. It also addressed the creation of a gold-standard corpus intended for training and evaluation of NLP methods. Future work will involve the development of a database to host and facilitate access to and analysis of information about mammalian milk oligosaccharides from scientific articles. Based on experimental results, a workflow will also be developed to automate the extraction of relevant information from large volumes of articles. For that purpose, the annotation schema and gold-standard corpus will serve as a basis for the standardized representation, extraction, and storage of knowledge.

## Acknowledgments

We are grateful to the INRAE MIGALE bioinformatics facility (MIGALE, INRAE, 2020. Migale bioinformatics Facility, doi: 10.15454/1.5572390655343293E12) for providing computing resources. The authors thank Agnès Girard (INRAE) for her help in bibliographic search on PuMed.

## Notes

### Competing Interest Statement

The authors have declared no competing interest.

## References

1. Walsh C, Lane J, van Sinderen D, Hickey R. Human milk oligosaccharides: Shaping the infant gut microbiota and supporting health. Journal of Functional Foods. 2020 Sep;72.

2. Ayechu-Muruzabal V, van Stigt A, Mank M, Willemsen L, Stahl B, Garssen J, et al. Diversity of Human Milk Oligosaccharides and Effects on Early Life Immune Development. Frontiers in Pediatrics. 2018 Sep 10;6.

3. Ruiz-Palacios GM, Cervantes LE, Ramos P, Chavez-Munguia B, Newburg DS. Campylobacter jejuni Binds Intestinal H(O) Antigen (Fucα1, 2Galβ1, 4GlcNAc), and Fucosyloligosaccharides of Human Milk Inhibit Its Binding and Infection. Journal of Biological Chemistry. 2003 Apr;278(16):14112–20.

4. Coppa GV, Zampini L, Galeazzi T, Facinelli B, Ferrante L, Capretti R, et al. Human Milk Oligosaccharides Inhibit the Adhesion to Caco-2 Cells of Diarrheal Pathogens: Escherichia coli, Vibrio cholerae, and Salmonella fyris. Pediatric Research. 2006 Mar;59(3):377–82.

5. Eiwegger T, Stahl B, Haidl P, Schmitt J, Boehm G, Dehlink E, et al. Prebiotic oligosaccharides: In vitro evidence for gastrointestinal epithelial transfer and immunomodulatory properties. Pediatric allergy and immunology. 2010 Dec;21(8):1179–88.

6. Bode L, Jantscher-Krenn E. Structure-Function Relationships of Human Milk Oligosaccharides. Advances in Nutrition. 2012 May 1;3(3):383S–391S.

7. Durham SD, Wei Z, Lemay DG, Lange MC, Barile D. Creation of a milk oligosaccharide database, MilkOligoDB, reveals common structural motifs and extensive diversity across mammals. Scientific Reports. 2023 Jun 26;13(1):10345.

8. Jin C, Lundstrøm J, Korhonen E, Luis AS, Bojar D. Breast Milk Oligosaccharides Contain Immunomodulatory Glucuronic Acid and LacdiNAc. Molecular & Cellular Proteomics. 2023 Sep;22(9):100635.

9. 9. Urashima T, Messer M, Oftedal OT. Oligosaccharides in the Milk of Other Mammals. In: Prebiotics and Probiotics in Human Milk. Elsevier; 2017. p. 45–139.

10. Thurl S, Munzert M, Henker J, Boehm G, Muller-Werner B, Jelinek J, et al. Variation of human milk oligosaccharides in relation to milk groups and lactational periods. British Journal of Nutrition. 2010 Nov;104(9):1261–71.

11. Trevisi P, Luise D, Won S, Salcedo J, Bertocchi M, Barile D, et al. Variations in porcine colostrum oligosaccharide composition between breeds and in association with sow maternal performance. Journal of Animal Science and Biotechnology. 2020 Mar 12;11(1).

12. Difilippo E, Willems HAM, Vendrig JC, Fink-Gremmels J, Gruppen H, Schols HA. Comparison of milk oligosaccharides pattern in colostrum of different horse breeds. Journal of agricultural and food chemistry. 2015 May 20;63(19):4805–14.

13. Samuel TM, Binia A, de Castro CA, Thakkar SK, Billeaud C, Agosti M, et al. Impact of maternal characteristics on human milk oligosaccharide composition over the first 4 months of lactation in a cohort of healthy European mothers. Scientific reports. 2019 Aug 13;9(1):11767.

14. Plows JF, Berger PK, Jones RB, Alderete TL, Yonemitsu C, Najera JA, et al. Longitudinal Changes in Human Milk Oligosaccharides (HMOs) Over the Course of 24 Months of Lactation. The Journal of nutrition. 2021 Apr 8;151(4):876–82.

15. Seferovic MD, Mohammad M, Pace RM, Engevik M, Versalovic J, Bode L, et al. Maternal diet alters human milk oligosaccharide composition with implications for the milk metagenome. Scientific reports. 2020 Dec 16;10(1):22092.

16. Azad MB, Robertson B, Atakora F, Becker AB, Subbarao P, Moraes TJ, et al. Human Milk Oligosaccharide Concentrations Are Associated with Multiple Fixed and Modifiable Maternal Characteristics, Environmental Factors, and Feeding Practices. The Journal of nutrition. 2018 Nov 1;148(11):1733–42.

17. van Leeuwen SS. Challenges and Pitfalls in Human Milk Oligosaccharide Analysis. Nutrients. 2019 Nov 6;11(11):2684.

18. Gopal PK, Gill HS. Oligosaccharides and glycoconjugates in bovine milk and colostrum. British Journal of Nutrition. 2000 Nov;84(S1):69–74.

19. Difilippo E, Pan F, Logtenberg M, Willems R (H. Am, Braber S, Fink-Gremmels J, et al. Milk Oligosaccharide Variation in Sow Milk and Milk Oligosaccharide Fermentation in Piglet Intestine. Journal of Agricultural and Food Chemistry. 2016 Mar 16;64(10):2087–93.

20. Remoroza CA, Liang Y, Mak TD, Mirokhin Y, Sheetlin SL, Yang X, et al. Increasing the Coverage of a Mass Spectral Library of Milk Oligosaccharides Using a Hybrid-Search-Based Bootstrapping Method and Milks from a Wide Variety of Mammals. Analytical Chemistry. 2020 Aug 4;92(15):10316–26.

21. Krallinger M, Rabal O, Leitner F, Vazquez M, Salgado D, Lu Z, et al. The CHEMDNER corpus of chemicals and drugs and its annotation principles. Journal of Cheminformatics. 2015 Dec;7(S1):S2.

22. Bada M, Eckert M, Evans D, Garcia K, Shipley K, Sitnikov D, et al. Concept annotation in the CRAFT corpus. BMC Bioinformatics. 2012 Dec;13(1):161.

23. Kim JD, Ohta T, Tateisi Y, Tsujii J. GENIA corpus--semantically annotated corpus for bio-textmining. Bioinformatics. 2003;19 Suppl 1:i180–182.

24. Kolařik C, Klinger R, Friedrich CM, Hofmann-Apitius M, Fluck J. Chemical Names: Terminological Resources and Corpora Annotation. Workshop on Building and evaluating resources for biomedical text mining (6th edition of the Language Resources and Evaluation Conference) Vol 36 2008.

25. Luoma J, Nastou K, Ohta T, Toivonen H, Pafilis E, Jensen LJ, et al. S1000: a better taxonomic name corpus for biomedical information extraction. Bioinformatics. 2023 Jun 1;39(6):btad369.

26. Bossy R, Deléger L, Chaix E, Ba M, Nédellec C. Bacteria Biotope at BioNLP Open Shared Tasks 2019. In: Jin-Dong K, Claire N, Robert B, Louise D, editors. Proceedings of the 5th Workshop on BioNLP Open Shared Tasks. Hong Kong, China: Association for Computational Linguistics; 2019. p. 121–31.

27. Albrecht S, Lane JA, Mariño K, Al Busadah KA, Carrington SD, Hickey RM, et al. A comparative study of free oligosaccharides in the milk of domestic animals. British Journal of Nutrition. 2014 Apr 14;111(7):1313–28.

28. Macias Rostami S, Bénet T, Spears J, Reynolds A, Satyaraj E, Sprenger N, et al. Milk oligosaccharides over time of lactation from different dog breeds. PLoS One. 2014;9(6):e99824.

29. Salcedo J, Frese SA, Mills DA, Barile D. Characterization of porcine milk oligosaccharides during early lactation and their relation to the fecal microbiome. Journal of dairy science. 2016 Oct;99(10):7733–43.

30. Urashima T, Saito T, Nakamura T, Messer M. Oligosaccharides of milk and colostrum in non-human mammals. Glycoconjugate Journal. 2001 May;18(5):357–71.

31. Chemical Entities of Biological Interest (ChEBI) [Internet]. [cited 2023 Jun 27]. Available from: https://www.ebi.ac.uk/chebi/

32. Schoch CL, Ciufo S, Domrachev M, Hotton CL, Kannan S, Khovanskaya R, et al. NCBI Taxonomy: a comprehensive update on curation, resources and tools. Database. 2020 Jan 1;2020:baaa062.

33. GeoNames [Internet]. [cited 2024 Sep 10]. Available from: http://www.geonames.org/

34. LBO | Summary | AgroPortal [Internet]. [cited 2024 Sep 10]. Available from: https://agroportal.lirmm.fr/ontologies/LBO

35. Rumeau M, Fenaille F, Girard A, Loux V, Ba M, Nédellec C, et al. MilkOligoThesaurus, a dataset of mammalian milk oligosaccharide synonyms. Data in Brief. 2024 Jun;54:110404.

36. Cejuela JM, McQuilton P, Ponting L, Marygold SJ, Stefancsik R, Millburn GH, et al. tagtog: interactive and text-mining-assisted annotation of gene mentions in PLOS full-text articles. Database. 2014 Apr 7;2014(0):bau033–bau033.

37. Papazian F, Bossy R, Nedellec C. AlvisAE: a collaborative Web text annotation editor for knowledge acquisition. Proceedings of the Sixth Linguistic Annotation Workshop 2012.

38. Lee J, Yoon W, Kim S, Kim D, Kim S, So CH, et al. BioBERT: a pre-trained biomedical language representation model for biomedical text mining. Bioinformatics. 2020 Feb 15;36(4):1234– 40.

39. Sung M, Jeon H, Lee J, Kang J. Biomedical Entity Representations with Synonym Marginalization. In: Proceedings of the 58th Annual Meeting of the Association for Computational Linguistics. Online: Association for Computational Linguistics; 2020. p. 3641–50.

40. Tizard IR. Comparative mammalian immunology: the evolution and diversity of the immune systems of mammals. Cambridge, MA: Academic Press; 2023.

41. Urashima T, Nakamura T, Yamaguchi K, Munakata J, Arai I, Saito T, et al. Chemical characterization of the oligosaccharides in milk of high Arctic harbour seal (Phoca vitulina vitulina). Comparative Biochemistry and Physiology Part A: Molecular & Integrative Physiology. 2003 Aug;135(4):549–63.

42. Tao N, Ochonicky KL, German JB, Donovan SM, Lebrilla CB. Structural Determination and Daily Variations of Porcine Milk Oligosaccharides. Journal of Agricultural and Food Chemistry. 2010 Apr 28;58(8):4653–9.

43. Xu G, Davis JC, Goonatilleke E, Smilowitz JT, German JB, Lebrilla CB. Absolute Quantitation of Human Milk Oligosaccharides Reveals Phenotypic Variations during Lactation. The Journal of Nutrition. 2017 Jan;147(1):117–24.

